# Scents Modulate Anxiety Levels, but Electroencephalographic and Electrocardiographic Assessments Could Diverge from Subjective Reports

**DOI:** 10.1101/2024.11.19.624243

**Authors:** Marina Morozova, Irina Gabrielyan, Daria Kleeva, Victoria Efimova, Mikhail Lebedev

## Abstract

Scents could modulate anxiety levels, such as anxiety in a medical office. Here we investigated the impact of two scents on the subjective and physiological anxiety markers in the dental office environment, utilizing self-reported anxiety assessments alongside physiological assessment with electroencephalographic (EEG) and electrocardiographic (ECG) measurements. Lavender was the first tested scent with the previously reported calming effect. African stone was the second stimulus with a musky scent. Twenty healthy participants took part in scent exposure sessions. Anxiety levels were assessed using the State-Trait Anxiety Inventory (STAI), EEG-based theta, alpha, and beta power ratios, and heart rate variability (HRV) indices derived from ECG data. Lavender exposure significantly decreased self-reported anxiety whereas African stone reduced physiological indicators of anxiety. Namely, African stone exposure led to decreased theta and increased alpha power in the parietal-occipital EEG signals. Additionally, decreases were observed in low-frequency (LF) HRV power and total HRV power, reflecting lowered autonomic arousal. These findings support the potential effectiveness for olfactory interventions to aid in anxiety management within clinical environments, but draw attention to the issue of proper evaluation of anxiety. In particular, the difference between the subjective reports and traditional EEG and HRV markers indicates that anxiety involves a complexity of factors, which makes its treatment by scents challenging.

## 1 Introduction

Anxiety is a frequent issue for patients in a medical office. Thus, visitors of a dental office view the clinical setting and tools and their anxiety grows as they anticipate discomfort and pain. Among the approaches to decreasing such anxiety, aromatherapy could be effective by acting on the emotions and their autonomic correlates. Anxiety-related stress responses are typically marked by heightened sympathetic nervous system activity and increased physiological arousal which could be monitored via electroencephalography (EEG) and electrocardiography (ECG) measurements. Here we investigated the effect of two scents on the anxiety levels of healthy participants placed in a dental environment. EEG and ECG recordings provided physiological data, and self-reported anxiety scores indicated how the participants experienced the scent effects subjectively.

### 1.1 EEG, ECG correlates of anxiety

EEG recordings offer insights into brain activity patterns associated with different emotional states, including anxiety and relaxation. EEG responses to anxious situations are often manifested as increased beta and theta power in the frontal cortex (Cavanagh and Shackman, 2014; Giannakakis et al., 2015), reduced asymmetry index (Muhammad and Al-Ahmadi, 2022) and reduced alpha power (Vanhollebeke et al., 2022). Increased beta and reduced alpha are also signs of heightened alertness often associated with anticipatory stress (Vanhollebeke et al., 2022). By examining changes in EEG patterns during scent exposure, it is possible to assess how specific odors affect neurological mechanisms of anxiety. Reductions in beta and increases in alpha activity typically accompany reduction in anxiety and better relaxation. Regarding the sensitivity of EEG to olfactory stimuli, previous studies have employed evoked potentials and EEG rhythms to analyze olfactory-induced responses (Sowndhararajan and Kim, 2016) and EEG correlates of such characteristics as odor pleasantness (Becerra et al., 2018) and attention to scents (Morozova et al., 2023). Furthermore, we have shown that odors could be used to provide neurofeedback of cortical activity (Medvedeva et al., 2024; Ninenko et al., 2024). Certain scents, such as lavender, have been shown to enhance alpha and theta activity (Sayorwan et al., 2012). The distinct effect of scents on EEG activity are potentially useful for the assessment of olfactory effects on the emotional states.

In addition to EEG, ECG provides a measure of the autonomic response to anxious situations. Anxiety is usually associated with increased heart rate and decreased heart rate variability (HRV) (Appelhans and Luecken, 2006; Chalmers et al., 2014; Kim et al., 2018). Previous studies have shown that soothing scents, such as lavender, increase the parasympathetic component of HRV and promote relaxation (Duan et al., 2006).

### 1.2 Scents in the Dental Office

Calming scents could be incorporated into dental offices to alleviate patient anxiety. Scents like lavender, orange and peppermint have been used in high-stress environments, such as medical and driving settings, with reductions in stress and fatigue (Czakert et al., 2024; Jiang et al., 2024; Lehrner et al., 2000).

In this study, we aimed to investigate the differential impacts of three distinct olfactory stimuli on anxiety in the dental office: water as a control, lavender oil and African stone, also referred to as hyraceum and noted for its musky scent.

Lavender oil was selected due to its prominence in the literature as a commonly studied aroma with calming effects, often associated with reductions in anxiety levels (Donelli et al., 2019). In contrast, African stone has controversial and understudied effects. African Stone is an animal essence used in perfumery. It is derived from the excrement of the rock hyrax, and the scent of hyraceum is described as animalistic with leather notes, similar to civet, musk, castoreum and ambergris. Hyraceum has been used in traditional South African medicine for various purposes, including treating epilepsy (Magama et al., 2018). Some studies have tested hyraceum samples for potential neuroactive properties, specifically affinity for GABA-benzodiazepine receptors (Olsen et al., 2007).

Through comparative analysis of these two scents, we aimed to elucidate the nuanced effects of the olfactory stimuli on anxiety, contributing to a deeper understanding of the role of scents in emotional regulation.

## 2 Materials and methods

### 2.1 Subjects

Twenty healthy volunteers — 10 females and 10 males, aged 36.0 ± 9.3 years (mean ± standard deviation), all right-handed — participated in the study. Each participant attended one experimental session lasting approximately 90 min. All participants gave informed consent, and the study received approval from the Ethics Committee of the Skolkovo Institute of Science and Technology, Moscow. Exclusion criteria included any history of neurological disorders or significant alterations in olfactory function within the previous six months.

### 2.2 Experimental setup

EEG data were collected using an NVX-36 amplifier (MKS, Russia). Recordings were made from 24 EEG channels following the international 10–20 system at a sampling rate of 500 Hz. Ag/AgCl electrodes with electrode gel were used, with a monopolar montage referenced to the FCz electrode. Electrode impedance was kept below 15 kΩ to ensure signal quality (Figure 1).

**Figure 1.**
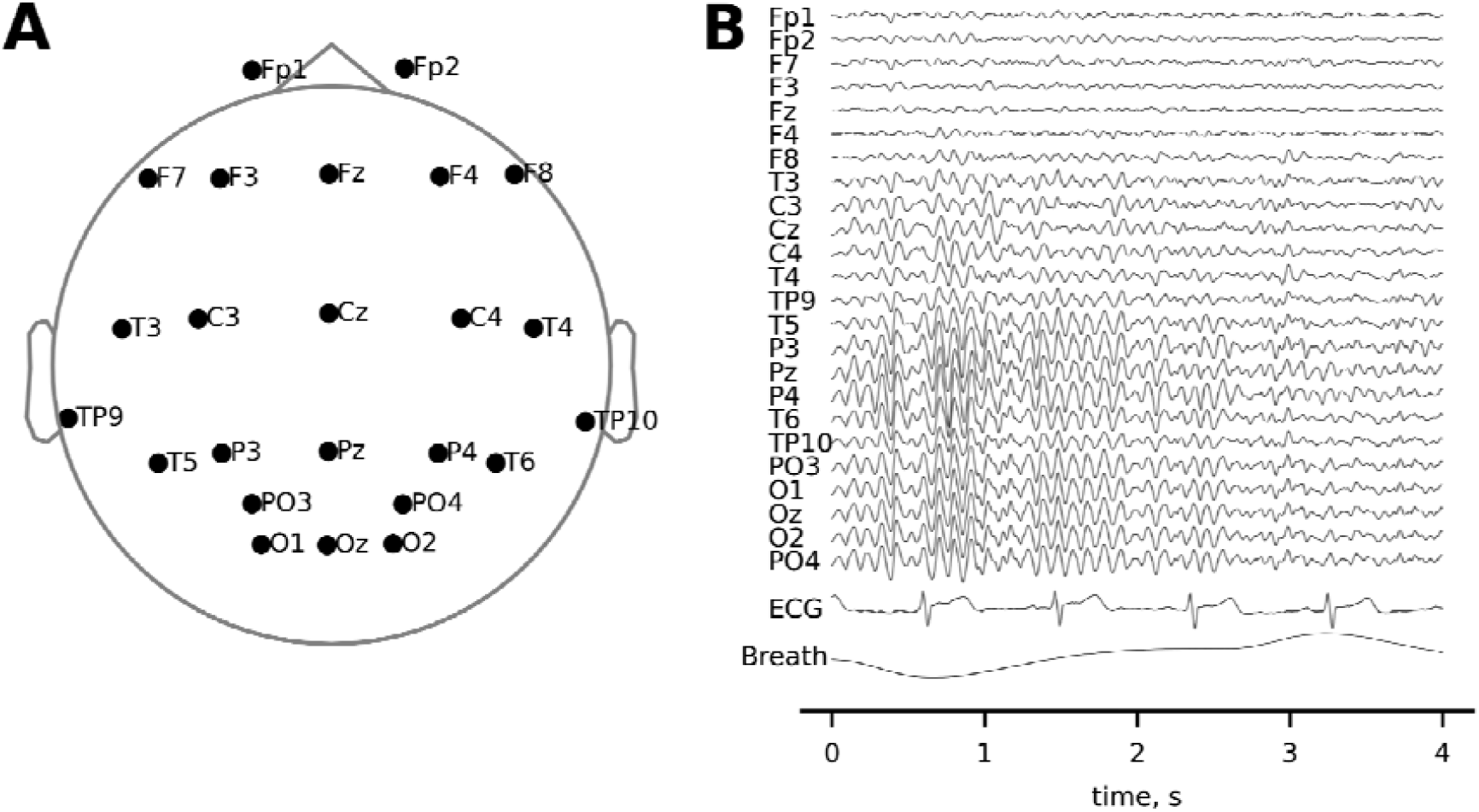
EEG settings. A: The layout of EEG channels. B: An example of the recorded EEG, ECG, and respiratory signals.

ECG data were collected through a separate channel connected to the left collarbone, and respiration was measured using a nasal thermometric breath sensor (TRSens, MKS, Russia).

### 2.3 Study design

The influence of scent exposure on anxiety levels were tested in a dental office. All participants reported some level of anxiety in these settings. Each participant was seated in a dental chair, and odorants were placed on a stand under the participant’s nose.

Three stimuli were tested: water (control), lavender oil, and African stone (hyraceum). Each stimulus was presented for 5 min in a randomized order, with participants inhaling the scent while their eyes were closed. During the intervals between the scent exposure sessions, the participants rested with their eyes open for 3 min, followed by 3 min with eyes closed. Two additional 3-min resting periods were recorded before the first and after the final scent presentation, allowing for baseline comparisons.

The full experimental protocol consisted of 11 recordings, including alternating periods of rest and scent inhalation:

1. Rest, eyes open (3 min)
2. Rest, eyes closed (3 min)
3. Inhalation of scent 1 (randomly selected: water, African stone, lavender), eyes closed (5 min)
4. Rest, eyes open (3 min)
5. Rest, eyes closed (3 min)
6. Inhalation of scent 2 (randomly selected), eyes closed (5 min)
7. Rest, eyes open (3 min)
8. Rest, eyes closed (3 min)
9. Inhalation of scent 3 (randomly selected), eyes closed (5 min).
10. Rest, eyes open (3 min)
11. Rest, eyes closed (3 min)

The participants completed the State-Trait Anxiety Inventory (STAI) prior to the experiment and immediately after each scent exposure, for a total of four STAI assessments. The STAI, a 20-item questionnaire, was used to measure both trait and state anxiety, with higher scores indicating greater anxiety.

### 2.4 Electrophysiological data analysis

Data preprocessing and analysis were conducted using Python libraries: MNE 1.3.1, SciPy 1.10.0, BioSPPy 2.1.2, and pyHRV 0.4.1.

To remove power line noise, a 50 Hz notch filter was applied to the EEG and ECG data. The EEG signals were band-pass filtered in the 1-40 Hz range using a 4th order Butterworth filter. The ECG data were high-pass filtered with a 0.1 Hz lower cutoff frequency using a 4th order Butterworth filter.

For EEG preprocessing, noisy channels were detected by visual inspection and interpolated using spherical spline interpolation. Eye-movement artifacts were removed with Independent Component Analysis (ICA), using Fp1-Fp2 channels as electrooculography (EOG) references.

Spectral analysis was performed using Welch’s method with a Hamming window. The EEG power spectra were computed across the 4-25 Hz range using a 2-s window and an 8-s FFT interval. Powers within theta (4-7 Hz), alpha (8-13 Hz), and beta (15-25 Hz) frequency ranges were calculated as ratios to the 4-25 Hz band for each electrode, providing data for comparison across conditions.

The ECG R-peaks were identified using the BioSPPy Python library (function biosppy.signals.ecg.ecg) and visually verified. Heart rate and NN-intervals were computed, with NN-intervals resampled at 4 Hz for spectral analysis using a 75-s length of each Welch segment and an 1024-second FFT length (function pyhrv.frequency_domain.welch_psd). Low frequency (LF, 0.04-0.15 Hz), high frequency (HF, 0.15-0.4 Hz), and total power (0-0.4 Hz) bands were calculated. LF and HF bands were normalized by dividing the power in each band by the total power in both LF and HF bands.

Respiratory cycles were detected (function biosppy.signals.resp.resp) and visually verified. Respiratory rate was calculated for comparison across the conditions.

### 2.5 Statistical analysis

The EEG ratio indices were analyzed with a permutation cluster-level paired t-test (function mne.stats.permutation_cluster_1samp_test) to compare theta, alpha, and beta ratios between scent inhalations for each electrode cluster, for a total of 9 permutation tests.

According to the permutation cluster-level paired t-test, clusters of parietal-occipital electrodes had statistically significant differences for the comparison of African stone with water, with theta ratios significantly lower and alpha ratios significantly higher during African stone exposure.

The parietal-occipital electrode theta and alpha ratios were averaged (across P3, Pz, P4, PO3, PO4, O1, Oz, O2 lead) for further paired comparisons across conditions.

Statistical comparisons of EEG indices, heart rates and respiratory rates across scent conditions were performed using non-parametric paired Wilcoxon tests.

LF, HF, and total HRV powers were baseline corrected by subtracting the values for the eye-open condition. The baseline-corrected values were compared across scent conditions using non-parametric paired Wilcoxon tests.

Scores from the STAI were analyzed using Wilcoxon tests to compare state anxiety scores across scent inhalations and baseline before the experiment, while Spearman’s rank correlation coefficient was used to examine the relationship between trait anxiety scores and physiological metrics.

## 3 Results

### 3.1 EEG findings

Statistical analysis of EEG data identified clusters of parietal-occipital leads where significant differences in theta and alpha power ratios were observed between inhalation of African stone and water (Figure 2). The theta ratio index had a significant decrease during African stone exposure compared to water (permutation cluster-level paired t-test, *p*-value = 0.036), with significant differences observed in leads C4, T4, T6, TP9, TP10, Pz, P4, PO3, PO4, O1, Oz, O2. Conversely, the alpha ratio index significantly increased during African stone inhalation compared to water (*p*-value = 0.043), with differences observed across parietal-occipital leads (P3, Pz, P4, PO3, PO4, O1, Oz). No statistically significant differences were found across conditions for the beta frequency range.

**Figure 2.**
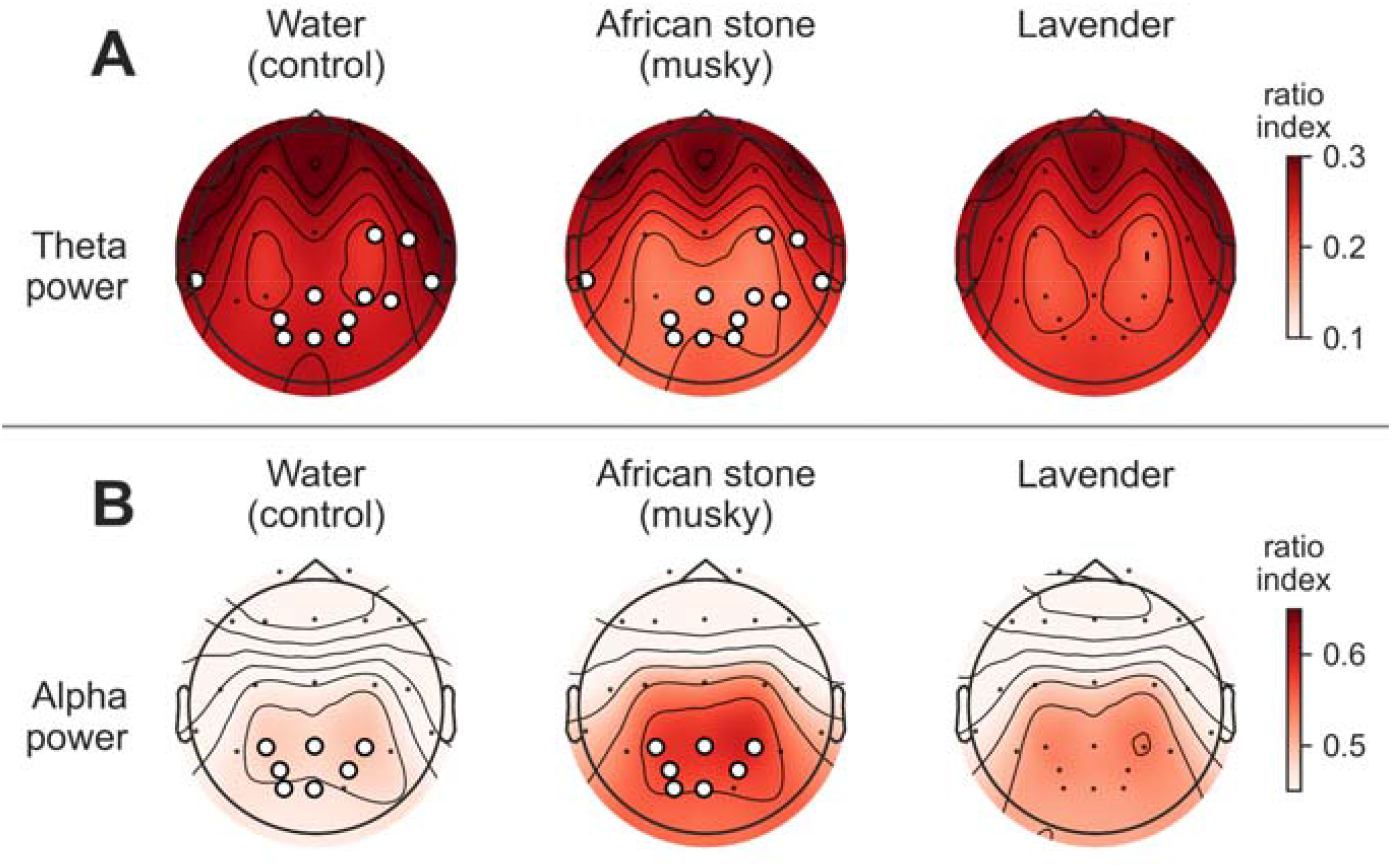
Topographic maps of mean EEG activity. A, B: theta and alpha ratio indices, respectively, across the conditions: water, African stone, and lavender. White dots indicate clusters of leads with statistically significant differences in the theta and alpha ratios between African stone and water.

In the subsequent analyses, theta and alpha ratios were averaged across the parietal-occipital leads (P3, Pz, P4, PO3, PO4, O1, Oz, O2) for comparisons across scent conditions. Using non-parametric paired Wilcoxon test, we confirmed a decrease in theta power during African stone inhalation relative to water (Figure 3A, *W* = 46, *p*-value = 0.027) and a significant increase in alpha power with African stone exposure (Figure 3B, *W* = 44, *p*-value = 0.021).

**Figure 3.**
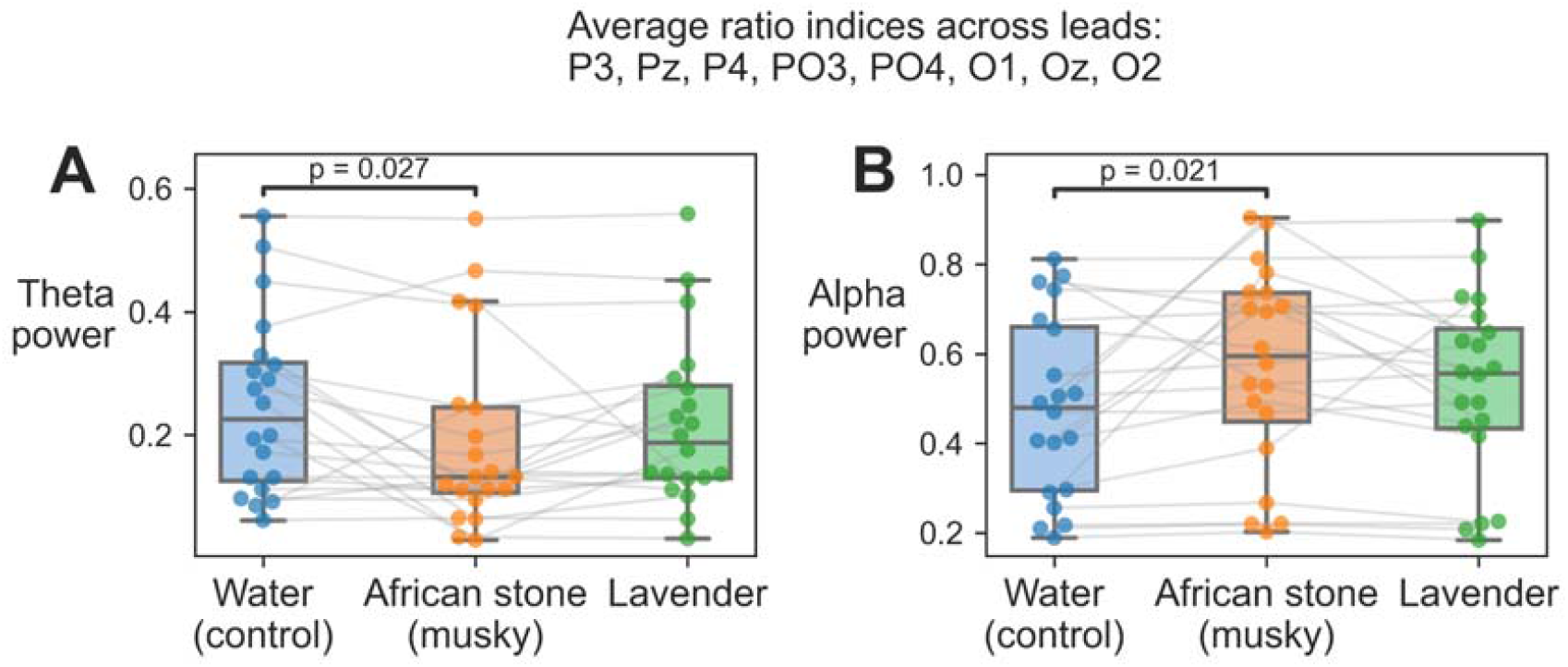
EEG power averaged for P3, Pz, P4, PO3, PO4, O1, Oz, and O2 leads. A, B: theta and alpha ratio indices, respectively for the comparison across conditions: water, African stone, and lavender.

### 3.2 HRV findings

No statistically significant differences were found in the heart rate and respiratory rate across the different scent conditions.

By contrast, baseline-corrected LF HRV power and total HRV power had significant reductions during African stone inhalation compared to lavender. Using a non-parametric paired Wilcoxon test, we found a significant decrease in the LF HRV power (*W* = 43, *p*-value = 0.019) and in total HRV power (*W* = 44, *p*-value = 0.021) during African stone inhalation relative to lavender (Figure 4).

**Figure 4.**
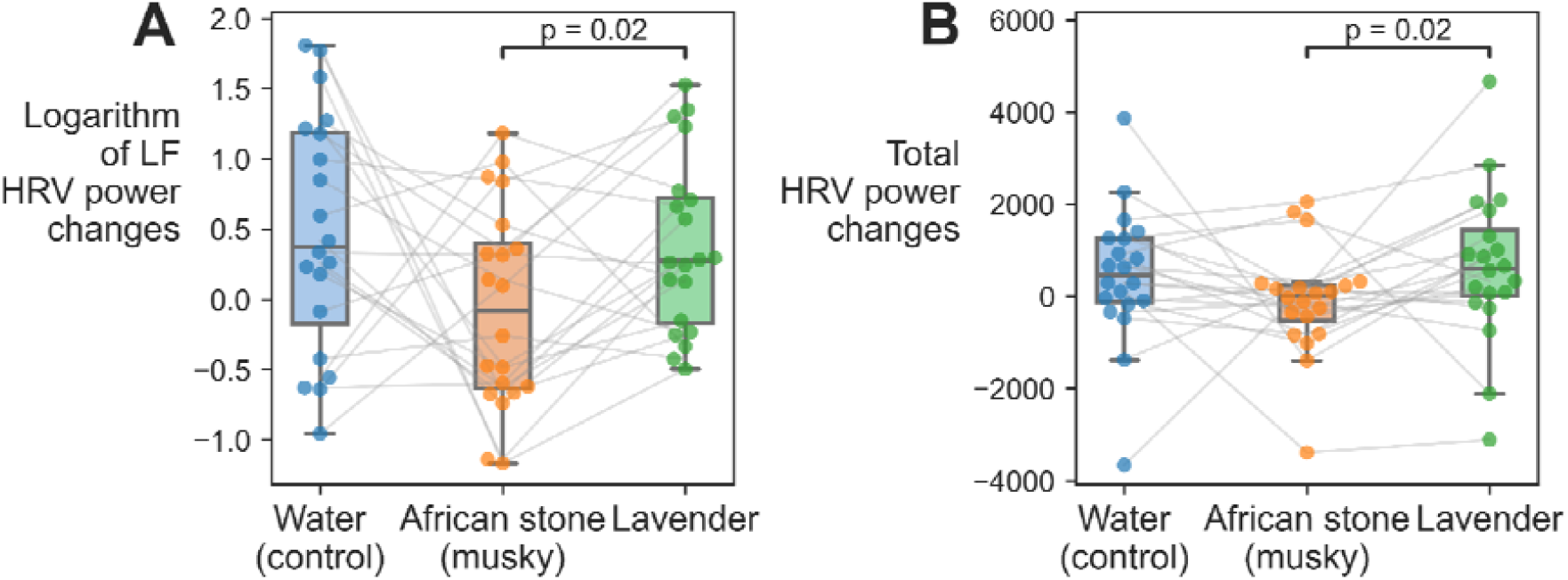
Changes in low frequency HRV (LF HRV, A) and total HRV power (B), relative to the preceding resting state with eyes open, across the conditions:water, African stone, and lavender.

### 3.3 Anxiety Assessment Findings

A significant reduction in state anxiety scores from the State-Trait Anxiety Inventory (STAI) was observed after lavender inhalation compared to the pre-experiment scores. Lavender inhalation significantly reduced the anxiety score (Figure 5A) compared to both pre-experiment (*W* = 38.5, *p*-value = 0.012) and water conditions (*W* = 42.5, *p*-value = 0.031). Following correction of anxiety scores against baseline (pre-experiment) scores, the reduction in state anxiety after lavender exposure remained statistically significant (Figure 5B, *W* = 42.5, *p*-value = 0.031).

**Figure 5.**
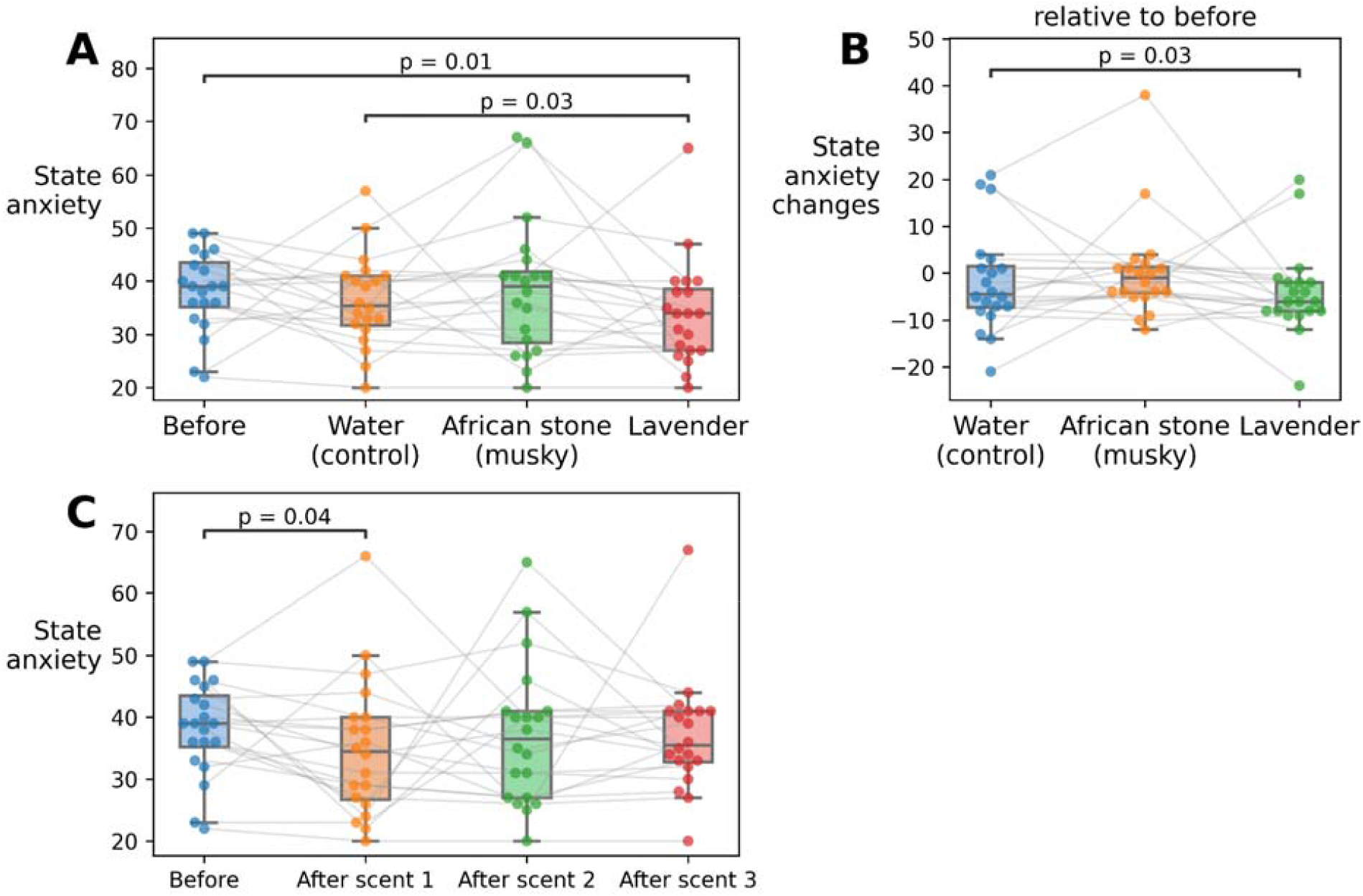
State anxiety scores from the STAI: A: The scores before the experiment and immediately after each odor inhalation, totaling four assessments. B) The baseline-corrected changes in state anxiety post-inhalation. C: The anxiety scores over the experimental session.

Notably, a general reduction in state anxiety scores was recorded over the course of the experimental session for all participants, with significant decreases after the first scent exposure compared to baseline (*W* = 50.5, *p*-value = 0.043) (Figure 5C).

### 3.4 Correlational Analysis

A moderate negative correlation was found between LF HRV power and trait anxiety scores from the STAI (Figure 6). Specifically, this was demonstrated by a negative correlation between the logarithm of LF HRV power and trait anxiety scores (Spearman’s rank correlation coefficient, *r*_*S*_ = -0.526, *p*-value = 0.0172), normalized LF HRV power and trait anxiety scores (*r*_*S*_ = -0.656, *p*-value = 0.0017), as well as between LF to HF ratio and trait anxiety scores (*r*_*S*_ = - 0.664, *p*-value = 0.0014). Additionally, there was a positive correlation between normalized HF HRV power and trait anxiety scores (*r*_*S*_ = 0.656, *p*-value = 0.0017), as normalized HF power is inversely related to normalized LF HRV power.

**Figure 6.**
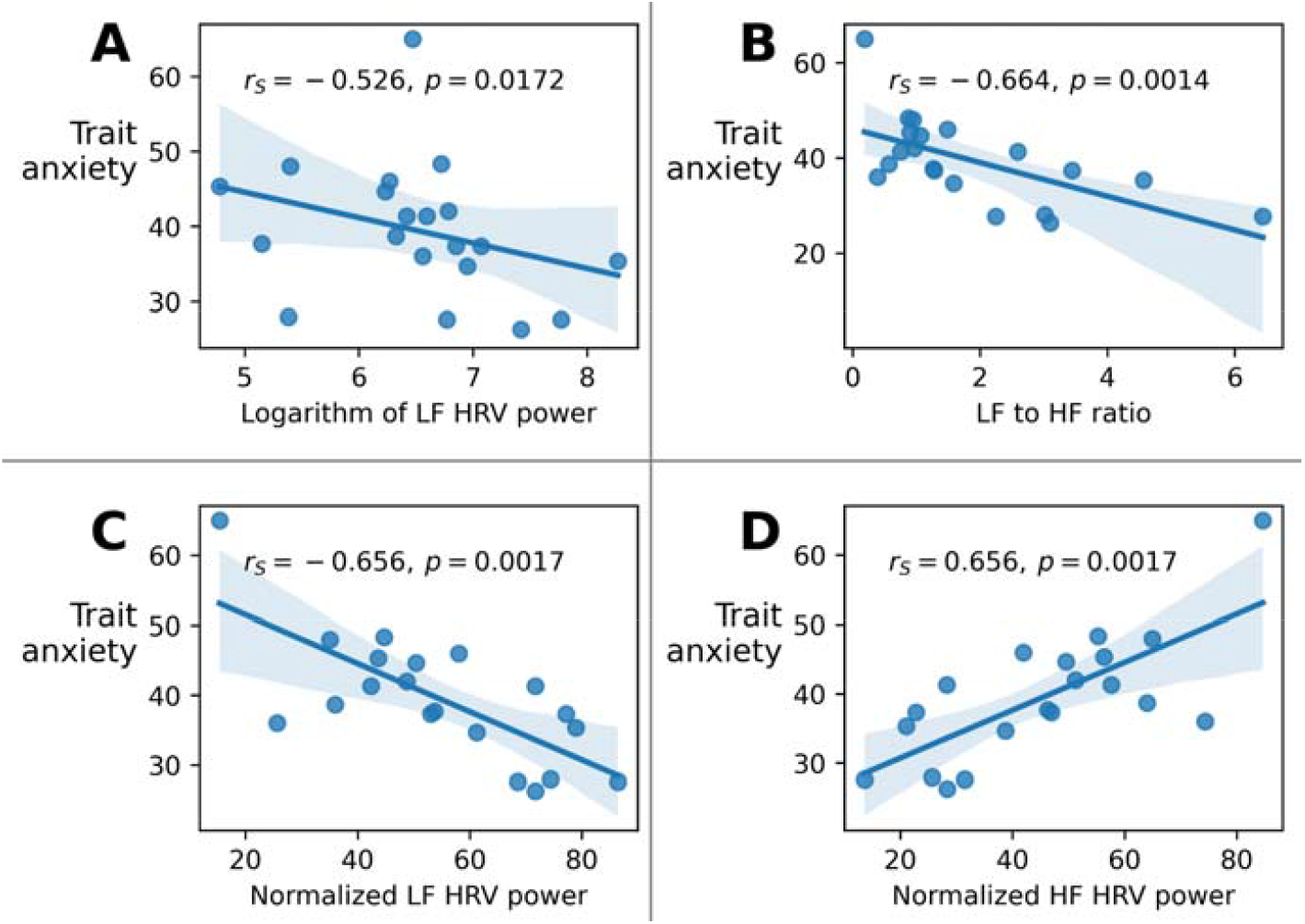
The relationship between trait anxiety scores and HRV metrics: A: Logarithm of low-frequency (LF) HRV power. B: LF to HF ratio. C: Normalized LF HRV power. D: Normalized HF HRV power.

## 4 Discussion

This study showed that lavender and African stone exhibited effects on anxiety-related metrics, with several differences between these scents. Consistent with prior findings, lavender inhalation notably decreased self-reported anxiety scores. This aligns with lavender’s established effect on subjective relaxation, potentially mediated by its influence on the limbic system and vagal tone reported in the previous studies of stress and autonomic regulation (Karan, 2019; Nagai et al., 2014; Sayorwan et al., 2012).

African stone had a stronger effect than lavender on the EEG metrics. It reduced EEG theta power and increased alpha power in parietal-occipital regions. Moreover, it decreased LF and total HRV power. These physiological changes could have resulted from a reduction in autonomic arousal common for the emotional state of relaxation and reduced anxiety.

Specifically, the decrease in LF HRV power and total HRV power during African stone inhalation could have been related to reduced sympathetic nervous system. These findings suggest that African stone could exert a calming physiological effect which is somewhat different from the subjective feeling of anxiety.

Such a divergence between the subjective and physiological markers of the scent effects on anxiety highlights the benefits of complex assessment of aromatherapy effects. Subjective assessment alone even could fail to capture physiological changes, particularly in the context of immediate or transient anxiety states. On the other hand, conventional EEG and ECG metrics could be influenced by a range of emotional and attentional states beyond anxiety, including general arousal or cognitive engagement, rather than short-term anxiety specifically. This limitation points to a broader issue in anxiety research: the potential need to reevaluate traditional anxiety biomarkers to better capture the specificity of short-term anxiety, especially in response to sensory interventions.

At this point in this research, we speculate that scent-based interventions could manage anxiety through different routes. Thus, lavender appears effective for reducing perceived anxiety whereas African stone appears to modulate physiological arousal. If our speculation is true, multifaceted aromatherapy could be utilized in clinical settings to exert specific and individual effects on the physiological arousal and subjective distress.

## 5 Conclusions

This study demonstrates that inhaling both lavender and African stone scents modulates anxiety-related metrics, but in distinct ways. Lavender is more effective for reducing self-reported anxiety, whereas African stone more strongly impacts physiological markers, including EEG theta and alpha power, and HRV parameters like LF and total power. This divergence between the effects of scenes on the subjective and physiological metrics underscores the need for comprehensive, multi-dimensional assessments of anxiety, as well as refinement in our understanding of EEG and HRV markers as reliable anxiety correlates, particularly in short-term and sensory-triggered contexts. These results support the potential for individualized olfactory interventions in clinical settings, where scents like lavender could be used for subjective comfort and African stone to target physiological relaxation.

## 6 Limitations

While this study provides valuable insights into the differential effects of lavender and African stone scents on subjective and physiological markers of anxiety, several limitations warrant consideration.

First, the sample size of 20 participants, while common in psychophysiological studies, limits the generalizability of these findings. Future research should aim to replicate these results in larger and more diverse populations to confirm the consistency of the effects observed here.

Second, the study’s reliance on EEG and HRV as primary physiological markers of anxiety introduces challenges in interpreting these data as definitive indicators of short-term state anxiety. EEG metrics, such as theta and alpha power, and HRV measures, including LF and total HRV power, reflect broader arousal and autonomic responses that may not be exclusively linked to anxiety. Further research with additional physiological markers (e.g., skin conductance, cortisol levels) could provide a more comprehensive understanding of the short-term effects of olfactory stimuli on anxiety.

Additionally, the divergence observed between subjective and physiological measures of anxiety highlights a potential limitation in using self-reported assessments to capture rapid or subtle shifts in anxiety state. The State-Trait Anxiety Inventory (STAI) and similar scales may not fully reflect the immediate impact of sensory interventions, such as scent exposure, especially in high-stress environments like dental offices. Future studies might benefit from using real-time, moment-to-moment anxiety tracking tools or exploring alternative self-assessment methods to capture transient effects more accurately.

Finally, individual differences in olfactory sensitivity, scent preferences, and baseline anxiety levels could have influenced participants’ responses to the scents. While measures were taken to control for some of these factors, these individual differences may still account for some variability in the results. Future research could address this by stratifying participants based on their olfactory sensitivity and scent preference profiles or by using standardized scent exposure levels.

By acknowledging these limitations, we hope to encourage further exploration of scent-based interventions in clinical settings, with an emphasis on refining both methodology and measures to enhance understanding and applicability in anxiety management.

## 8 Author Contributions

MM contributed to constructing the experimental setup and data analysis. IG contributed to constructing the experimental setup and data collection. MM, IG, VE, DK and ML equally contributed to the development of the experiment design and writing of the manuscript. All authors contributed to the article and approved the submitted version.

## 10 Data Availability Statement

The raw data supporting the conclusions of this article will be made available by the authors, without undue reservation.

